# GSNOR deletion differentially alters age-related cardiac function in a sex-dependent manner

**DOI:** 10.1101/2025.07.28.667309

**Authors:** Obialunanma V. Ebenebe, Raihan Kabir, Allison Booher, Haley Garbus, Charles D. Cohen, Vivek Jani, Brian L. Lin, Luigi Adamo, Mark Kohr

## Abstract

S-nitrosoglutathione reductase (GSNOR), a regulator of protein S-nitrosylation (SNO), has been proposed as a longevity protein. GSNOR signaling has been implicated in both the alleviation and exacerbation of aging. In the context of ischemia reperfusion injury, we previously showed a sex-dependent response to GSNOR inhibition; cardiac damage was alleviated in males and exacerbated in females. Considering sex differences in the incidence of cardiovascular disease with age, we investigated the effect of GSNOR deletion (^-/-^) on age-related changes in cardiac function. We performed longitudinal 2D-echocardiography measurements in M-Mode on male and female, wildtype (WT) and GSNOR^-/-^ mice at young (3-4 months), middle (13-15 months) and old age (18-20 months). Left ventricular wall thickness and ejection fraction decreased with age in WT mice but was maintained in GSNOR^-/-^. Western blot and GSNOR-activity assay showed GSNOR activity and expression decreased with age in WT females alone. Isolated cardiomyocyte force-coupling analysis showed increasing age was inversely correlated with sarcomere shortening and Ca^2+^ release kinetics in WT males, but not GSNOR^-/-^. WT females showed slower Ca^2+^ re-uptake after contraction and time to peak sarcomere shortening, but all other parameters were maintained. GSNOR^-/-^ females exhibited slower Ca^2+^ re-uptake and decreased sarcomere shortening. Proteomic analysis of SNO from females showed upregulation of Pyruvate Dehydrogenase, E1 Beta and Dihydrolipoamide dehydrogenase in young WT females relative to middle-age mice. Together our data suggest that GSNOR deletion is cardioprotective by maintaining cardiac function in males; while in females the absence of GSNOR removes an age-essential SNO-imbalance, which may exacerbate pathologies.

## INTRODUCTION

Cardiovascular disease (CVD) continues to be a leading cause of mortality in men and women in the US [1], yet mechanisms underlying age-related increases in adverse CVD events are still unknown. Furthermore, the incidence of CVD is sex-dependent, with women having a two-fold higher likelihood of developing CVD later in life and higher CVD related mortality relative to age-matched men [2, 3].

Proposed mechanisms for the observed sex differences include nitric oxide (NO) signaling and associated protein *S*-nitrosylation (SNO), which are thought to drive female-specific cardioprotective mechanisms involved in reducing cardiac damage during ischemia-reperfusion (I/R) injury [4–7]. Cellular levels of protein SNO are governed in part by the enzyme *S*- nitrosoglutathione reductase (GSNOR). Previous studies, including those from our group [8] and others [9, 10], have shown a sex-dependent response to GSNOR inhibition in the context of I/R injury in mice. These murine models demonstrated that the loss of GSNOR alleviated cardiac damage in male mice, but injury was exacerbated in female mice. GSNOR has also been implicated in age-related maladaptation in other organs [11] and cellular models [12]. Collectively, these findings suggest that GSNOR may play a critical role in sex-dependent aging-induced CVD outcomes.

In the current study, we explored basic cardiac and morphometric changes in wildtype (WT) and GSNOR knockout (KO) mice during aging in isolated cardiomyocytes and at the level of the whole heart . Following these mice from young to old age, we observed sex-dependent and age-related changes in heart weight, individual cardiomyocyte function and overall cardiac phenotype. Our data show that the absence of GSNOR alters “normal” cardiac aging in our mouse model. This study aimed to showcase a role for GSNOR in age- and sex-dependent CVD changes with a particular emphasis on understanding the heightened risk in aging females.

## MATERIALS AND METHODS

### Animals

All procedures were approved by the Johns Hopkins University Institutional Animal Care and Use Committee and conducted in accordance with the Guide for the Care and Use of Laboratory Animals published but the United Stated National Institute of Health (NIH publication No. 85±23, revised 2011). Generation of GSNOR^-/-^ mice (C57Bl/6J background) was previously described [8]. All male and female wild-type (WT) C57Bl/6J mice used in the study were purchased from Jackson Laboratories (Bar Harbor, ME). For repeated procedures, eleven-week-old WT mice were purchased and aged to appropriate time points; for isolated procedures, WT mice were purchased and age and sex-matched to GSNOR^-/-^ mice. All mice were housed (4-5 per cage) under specific pathogen-free conditions at Johns Hopkins University. Mice were maintained on standard chow and water *ad libitum*. Mice described as “Young” are 3-4 months old, mice described as “Middle-age” are 13-15 months old, and mice described as “Old” are 18-20 months old.

### Echocardiography

Transthoracic echocardiography measurements were performed and analyzed by technicians blinded to the experimental groups at the Johns Hopkins Small Animal Cardiovascular Phenotyping and Model Core, as previously described [8, 13, 14]. Echo measurements were carried out on conscious mice using a Vevo 2100 system with a 40-MHz linear transducer (FUJIFILM VisualSonics, Toronto, Canada). The M-mode echocardiogram was acquired from the short axis view of the left ventricle (LV) at the level of the midpapillary muscles, by an independent read. From this axis view of the left ventricle, the following cardiac parameters were measured: inter-ventricular septal thickness at end-diastole (IVSd) and end-systole (IVSs), LV internal diameter at end-diastole (LVIDd) and end-systole (LVIDs), LV posterior wall thickness at end-diastole (LVPWd) and end-systole (LVPWs), ejection fraction (EF) and fractional shortening (FS). Using these parameters, left ventricle relative wall thickness (RWT), cardiac output (CO) and left ventricle mass (LV mass) were calculated in the estimation of cardiac contractility and left ventricle morphology. These indices were derived from the following equations:

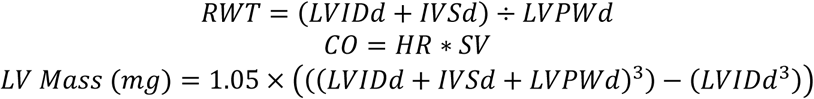

### Heart Perfusion

For all procedures, mice were anesthetized with a mixture of ketamine (90mg/kg; Hofspira Inc., Lake Forest, IL) and xylazine (10mg/kg; Sigma, St. Louis, MO) via intraperitoneal injection, and anticoagulated with heparin (Fresenvis Kabi, Lake Zurich, IL). After verifying adequate anesthesia via toe pinch, mice were subsequently euthanized via myocardial excision and exsanguination. Hearts were rapidly excised via thoracotomy and arrested in ice-cold Krebs-Henseleit buffer. Excess tissue and fat were trimmed, and hearts were cannulated onto a Langendorff apparatus, and retrogradely perfused (5 minutes) under constant pressure (100 cmH_2_0) and temperature (37°C), with oxygenated Krebs-Henseleit buffer (95 % O_2_, 5 % CO_2_; pH 7.4) to clear coronary vessels of blood. After perfusion hearts were weighed, sectioned into two halves and snap-frozen with liquid nitrogen.

### Heart homogenate preparations

All procedures were done in the dark to prevent light-induced cleavage of protein SNO residues. Hearts were homogenized in assay-specific buffers using a Precellys/Cryolys Evolution Homogenizer (3 × 30 s cycles, 20s pause, 0 ◦C, 7200 RPM; Bertin Instruments), with a hard tissue homogenizing kit (CK28) as previously described [14]. Homogenates were incubated on ice for 15-20 minutes, then ultracentrifuged for 10 minutes at 14000 *g* at 4°C. Supernatant was collected as crude heart homogenate into amber tubes; protein concentration was determined using the Bradford Assay. Homogenates were aliquoted and stored at -80°C until further analysis.

### GSNOR Activity Assay

GSNOR activity was determined from samples homogenized in cell lysis buffer (Cell Signaling Technology, Danvers, MA) and supplemented with a Protease/Phosphatase Inhibitor Cocktail (Cell Signaling Technology), as previously described [8, 13]. Briefly heart homogenates (100μg) were diluted in assay buffer consisting of (in mM): Tris-HCl (20), EDTA (0.5), Neocuproine (0.5), 1% Tergitol and Protease/Phosphatase Inhibitor Cocktail (Cell Signaling Technology). Diluted samples were loaded into individual wells on a 96-well plate on ice. NADH was made fresh before use with assay buffer and added to each sample well (final concentration-200μM). Basal NADH consumption was recorded at absorbance wavelength 340nm for five minutes at 25°C. GSNO was then added to each sample to initiate reaction and NADH consumption was recorded for 30 minutes. The deviation of the regression slope from zero was used to extrapolate GSNOR activity.

### GSNOR Protein Expression by Western Blot

Heart homogenates (30µg) were separated (20 minutes, 75V; 80 minutes, 150V) on a gradient Bis-Tris SDS-PAGE gel (4-12%, NuPAGE; Thermo Fisher, Carlsbad, CA) and transferred (90 minutes, 220mA) to a Polyvinylidene fluoride membrane. Every gel included a molecular weight marker to identify the region of interest (Novex Prestained Protein Standard, Thermo Fisher). Covalently labelled lysine residues of the total protein, visualized by fluorescence imaging (488nm), served as loading control (No-Stain Protein Labelling reagent, Thermo Fisher) following manufacturer’s instructions. Membranes were blocked (1 hour) at room temperature (rt) with bovine serum albumin (5% wt./vol; Sigma-Aldrich) in Tris-buffered saline with 0.1% Tween-20 (TBS-T) and subsequently incubated overnight (4°C) with primary antibody against GSNOR (1:1000, abcam:175402). Membranes were washed with TBS-T (30 minutes) prior to secondary antibody incubation (1 hour, rt) (1:5000, anti-rabbit-HRP, Cell Signaling Technology: 7074S) and visualized using chemiluminescence substrate (SuperSignal West Pico PLUS, Thermo Fisher) on an iBright Imager (Thermo Fisher).

### Cardiomyocyte isolation

Myocytes were isolated as previously described [15, 16]. Briefly, mice were anesthetized with isoflurane and subsequently euthanized via cervical dislocation. Hearts were rapidly excised via thoracotomy and retrogradely perfused via the aorta at a constant flowrate of 1.5mL/min with a Ca^2+^-free buffer containing (in mM): glucose (5.5), taurine (5), MgCl_2_ (1), butanedione monoxime (5). This was followed by enzymatic digestion with collagenase type II (Worthington) and protease type XIV (Sigma) for a total perfusion time of 9-10 minutes. Hearts were then mechanically digested, and triturated gently with a pipette. Dispersed cardiomyocytes were then filtered and centrifuged at 800rpm for 1 minute. The cardiomyocyte pellet was resuspended in a digestion stop solution of Tyrode buffer supplemented with 0.4% (w/v) Bovine Serum Albumin (BSA) containing (in mM): NaCl (1.4), KCl (5) HEPES (10) CaCl_2_ (0.125). Myocytes were subsequently transferred to BSA-free Tyrode solution and the calcium concentration was gradually increased every 10 minutes to a final concentration of 1mM. Only live cardiomyocytes were used for measurements. Prior to measurements, cardiomyocytes were incubated in 1M Fura-2AM for 10- 14 minutes and then washed with fresh 1mM Ca^2+^-Tyrode buffer. Cardiomyocytes were used for functional measurements within 5 hours of isolation. Ca^2+^ transients and sarcomere shortening were measured using an inverted microscope (Nikon Eclipse TE-2000U) and a custom-built IonOptix system with IonWizard software as previously described [17]. Cardiomyocytes were stimulated at 1 Hz and maintained at 37°C in Tyrode buffer for all measurements. Fura-2-AM was excited at 340 and 380 nm alternating at 250 Hz and emission recorded at 510 nm by a single photon multiplier tube. Ca^2+^ signal transients (10-15 measurements), along with corresponding sarcomere shortening, were averaged per cardiomyocyte, and a minimum of 10 cardiomyocytes per mouse were included in analyses. The fluorescence ratio value is shown as F/F_0_ (fluorescence normalized to baseline 340/360).

### SNO-Resin Assisted Capture (SNO-RAC) with Liquid Chromatography and Mass Spectrometry Sample preparation

SNO sites were identified using a SNO-RAC protocol previously described [18, 19], from hearts homogenized in HEN buffer containing (in mM, ph7.7): sucrose (300), **H**EPES-NaOH (250), **E**DTA (1), **N**eocuproine (0.1), 0.5% Triton-X 100 and N-ethylmaleimide (25; NEM, Sigma), supplemented with an EDTA-free protease inhibitor tablet (Roche). All procedures were performed in the dark and buffers were degassed before use to prevent light-cleavage of SNO residues and oxidation of the resin, respectively. Briefly, samples (2mg protein) were diluted in HEN buffer supplemented with 20mM NEM, oscillated at 800 rpm at 50°C for 40 minutes to block unmodified free thiol groups. Excess NEM was removed using Zeba desalting columns (Thermo Fisher, Waltham, MA). Thiol-binding thiopropyl sepharose resin was rehydrated for 25mins with HPLC-grade water. Following rehydration, 200µL of the resin slurry was used per sample in Pierce Mini Spin Columns (ThermoFisher). Samples were incubated in the resin with 30mM ascorbate, and oscillated in the dark for 3 hours at 37°C. Resin-bound proteins were washed with HENS buffer and diluted (1:10) in HENS buffer to remove SDS. Samples were then trypsin digested (Promega, WI) in 10mM ammonium bicarbonate overnight with oscillation at 37°C. To further reduce SDS and salts, resin-bound peptides were washed with diluted HENS buffer, NaCl and HPLC-grade water before elution at room temperature for 30mins in 20mM dithiothreitol and 10mM ammonium bicarbonate. Eluted peptides were alkylated with 40mM d5-NEM at 37°C for 40mins, acidified with 0.1%TFA and cleaned using 10µg C18 zip-tips. Peptides were eluted with 60% acetonitrile and 0.1% TFA, dried using a Savant SpeedVac (Thermo), and stored in -20°C.

### Liquid chromatography separation and tandem mass spectrometry (LCMS/MS)

Dried peptides were reconstituted in 2% ACN and 0.1% FA and analyzed by nanoflow LC-MS/MS using a Neo Vanquish UHPLC interfaced with an Orbitrap Exploris 480 mass spectrometer (Thermo Fisher Scientific). Peptide separation was performed with a linear gradient (water/acetonitrile) over a polyimide-coated, fused-silica, 25 cm × 360 μm o.d./75 μm i.d. self-packed PicoFrit column (New Objective) with a built-in emitter (75 µm emitter i.d.). Stationary phase in the analytical column consisted of ReproSil-Pur 120 C18-AQ, 2.4 μm particle size, 120 Å pore (Dr. Maisch High Performance LC GmbH). The trap column consisted of ∼1 cm × 360 μm o.d./75 μm i.d. polyimide-coated, fused-silica tubing (New Objective), packed with 5 μm particle size, 120 Å pore, C18 stationary phase (ReproSil-Pur), with a Kasil frit. Electrospray ionization was accomplished with 2 kV positive spray voltage and an ion transfer tube temperature of 250 °C. For data-dependent acquisition (DDA) label-free analysis, peptides were separated by a 100- minute linear gradient. Survey scans (MS1) of precursor ions were acquired in the Orbitrap detector of the mass spectrometer from 350-1800 m/z at 120,000 resolution at 200 m/z, automatic gain control (AGC) set to standard limits, injection time set to Auto, and an RF lens setting of 50%. The top 15 precursor ions from each survey scan were isolated with a 1.5 m/z window, a 15s dynamic exclusion (with a ±10 ppm tolerance), and the following filters: intensity threshold of 5x104, charge states 2-6. Precursors were fragmented by HCD with a normalized collision energy of 30% and MS/MS spectra were acquired at 30,000 resolutions, AGC set to standard limits, and injection time set to Auto.

### Peptide quantification and database search

All database searches were conducted using Proteome Discoverer v3.1 (ThermoFisher Scientific) with Mascot (version 2.8.3). Data was searched against a mouse UniProt FASTA database (proteome accession UP000000589, 63785 entries) and a custom database containing common contaminants (e.g., human keratin; 438 entries). Search criteria were tryptic cleavage (maximum 2 missed), 15 ppm precursor and fragment ion mass tolerances, and variable modifications of Cys carbamidomethylation, Met oxidation, Asn/Gln deamidation, Cys nitrosylation, and Cys modification with D5-N-ethylmaleimide. Peptide identifications were validated by Percolator at 1% false discovery rate (FDR) based on an auto-concatenated decoy database search and modification sites were localized with IMP-ptmRS. Label-free quantitation was performed using MS1 peak areas of unique peptides and values were normalized to total peptide abundances. Average normalized abundances were then used to calculate fold change (log2FC) and statistical differences between groups[20].

### Statistical analysis

Data are presented as mean ± SEM. Statistical significance (p<0.05) was determined as appropriate by linear regression or ANOVA, followed by a Tukey’s post hoc test for multiple group comparisons. Data processing and analyses were done using Excel (Microsoft 365), GraphPad Prism (version 10.4.) and RStudio (R version 4.3.3).

## RESULTS

### Cardiac GSNOR expression and activity is changed in a sex-dependent manner with age

We first examined how GSNOR activity is altered in the heart with aging. To this end we measured GSNOR activity, determined by the rate of NADH consumption in the presence of GSNO, in hearts from young and middle-age WT male and female mice. The regression slope was unchanged between young and middle-age males (Figure 1A), but there was a significantly greater elevation in middle-age male hearts relative to young males (p<0.0001). This suggests that although GSNOR activity in males is unchanged, there may be additional NADH-consuming enzymes active in young male hearts relative to middle-age males. Interestingly in females, there was a statistically significant increase in the regression slope of the middle-age group relative to young (p=0.05; Figure 1B), suggesting GSNOR activity decreases in middle-age female hearts. Due to the difference between the slopes, it was not possible to test whether the intercepts differ significantly. We then looked to see whether this difference in activity was due to differences in GSNOR protein expression. Here we included an old group to fully demarcate the aging effect. In male mice, cardiac GSNOR protein expression levels were unaffected by age (Figure 2A). Interestingly in female hearts, there was no change in GSNOR protein expression from young to middle-age, but there was a significant decrease in GSNOR expression in old female hearts relative to young and middle-age mice (p=0.0061 and 0.0089, respectively; Figure 2B). Collectively, our findings demonstrate that GSNOR expression is maintained in young to middle-aged females, but GSNOR activity decreases.

**Figure 1:**
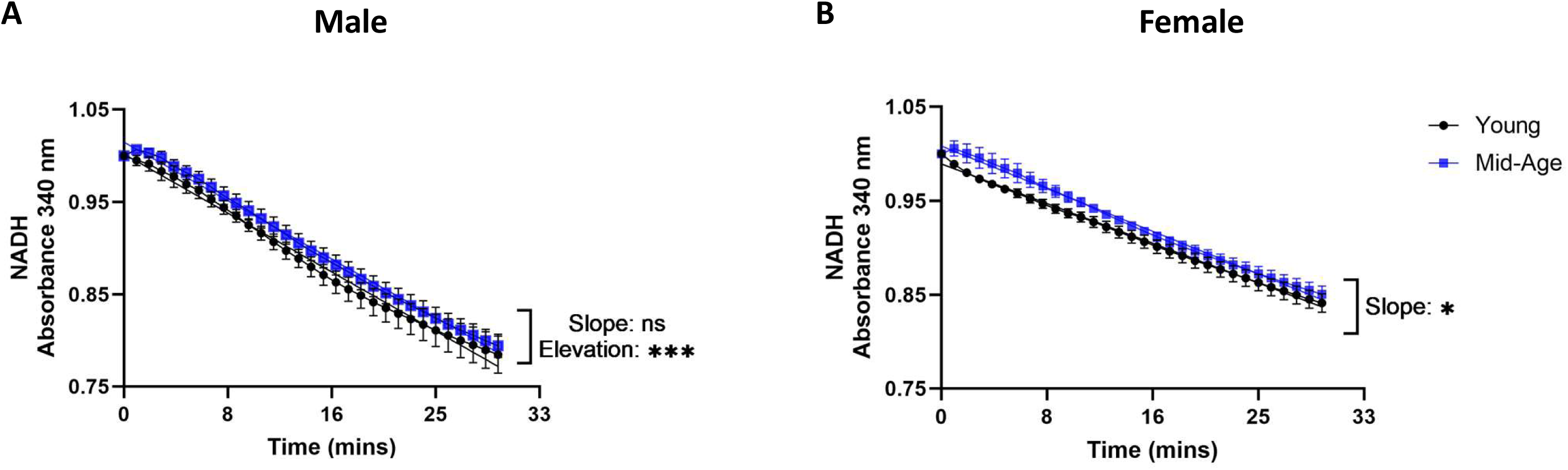
S-nitrosoglutathione reductase (GSNOR) activity in wild-type mice. GSNOR activity in heart homogenates from young (3-4 months; black) and mid-age (13-15 months; blue) male (**A**) and female (**B**) mice measured by NADH absorption in the presence of GSNO over time. Data are normalized to reading at Time-0. Data presented mean ± SEM of 2 technical replicates; n=5/group. Simple linear regression to determine difference between slopes, elevation and intercept. Statistical significance was determined at p<0.05.

**Figure 2:**
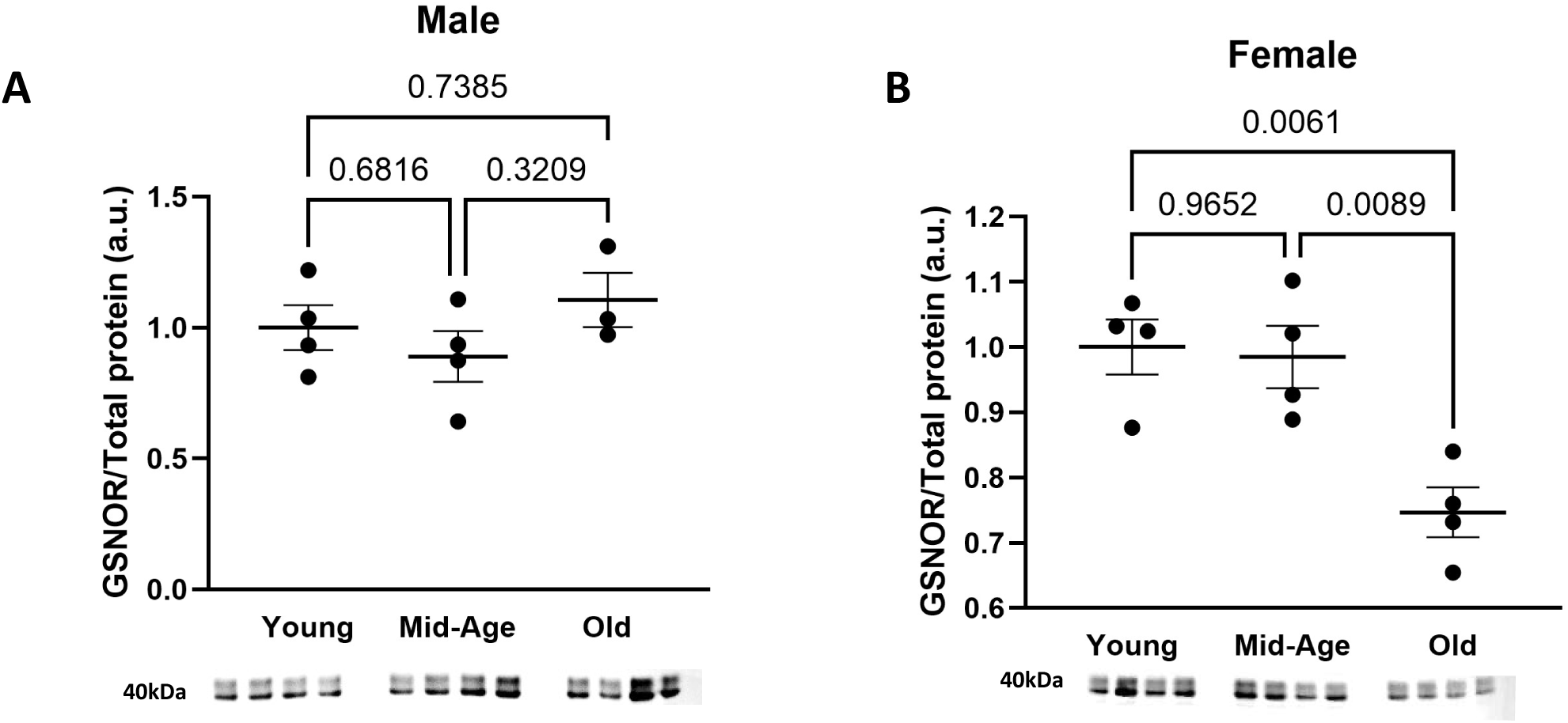
S-nitrosoglutathione reductase (GSNOR) expression in wild-type mice. Densitometry quantification of GSNOR expression in heart homogenates from young (3-4 months), mid-age (13-15 months) and old (18-20 months) male (**A**) and female (**B**) mice. Data are normalized to mean of young group and are presented ± SEM; n=4/group. Statistical significance was determined at p<0.05 by Ordinary one-way Anova with Tukey’s post-hoc test.

### GSNOR deletion alters age-related changes in heart weight

As a next step, we assessed basic morphometric changes with age in WT mice and compared them to age-matched GSNOR^-/-^ mice. As previously documented [21], there was a significant effect of age on body weight (BW) in both sexes of WT and GSNOR^-/-^ mice (p<0.0001; Figure 3). In both sexes of WT mice, there was a significant increase in BW comparing young to middle-age (p<0.0001; Figures 3A & 3B) and young to old (p<0.0001; Figures 3A & 3B) groups. There was no overt change in BW of both male and female WT mice from the middle-age to the old group. However, in GSNOR^-/-^ mice, while BW followed a pattern of increase similar to WT mice this was not to the same extent in both sexes. BW was increased in male GSNOR^-/-^ mice from young to middle-age and young to old age (p=0.025 and p=0.003 respectively). At all age groups, male

**Figure 3.**
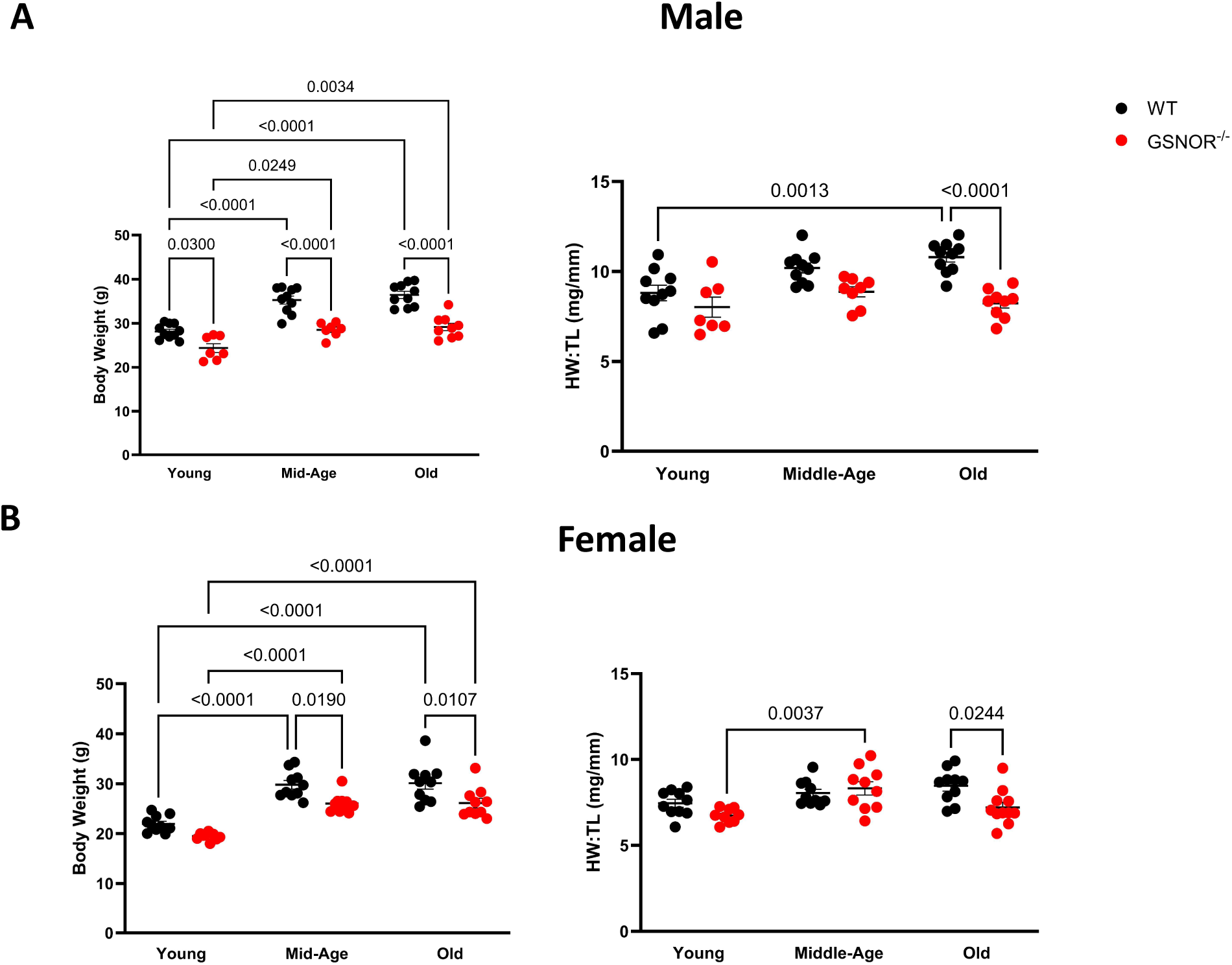
Body and heart weight measurements in wild-type and GSNOR-KO mice. Young (3-4 months), mid-age (13-15 months) and old (>18 months) male (**A**) and female (**B**) wild-type (WT; black) and GSNOR-KO (red) mice were weighed following anesthesia, prior to termination and heart excision. Heart weights (HW) were obtained following 5-mins perfusion of KRH buffer on Langendorff system and normalized to body weight (BW). Data represents mean ± SEM. Statistical significance was determined by two-way ANOVA with Tukey’s post-hoc test; n=10-15 mice/group.

GSNOR^-/-^ mice were significantly smaller than their WT counterparts (Figure 3A). In female GSNOR^-/-^ mice a significant BW increase was observed between young to middle age (p<0.0001; Figure 3B) and young to old (p<0.0001; Figure 3B). Interestingly, relative to their WT counterparts female GSNOR^-/-^ BW was the same in the young group; slight decreases in BW were observed in the middle-age and old groups of female GSNOR^-/-^ mice relative to WT (p=0.02 and p=0.01, respectively; Figure 3B).

Similar to previous reports [21, 22], we observed an increase in heart weight (HW) normalized to tibia length with age in WT mice. In male WT mice, HW was significantly greater in old relative to young mice (p=0.0013; Figure 3A). Female WT mice showed a similar pattern of HW increase with age, but not as robust as the males, once normalized to tibia length (Figure 3B). In male GSNOR^-/-^ mice, the HW increase with age was modest; hearts from old GSNOR^-/-^ male mice were significantly smaller than their WT counterparts (p<0.0001; Figure 3A).

Female GSNOR^-/-^ mice showed a significant increase in HW from young to middle-age group (p=0.004; Figure 3B); this HW increase in the middle-age GSNOR^-/-^ mice seemed comparable to WT. Interestingly, there was also a decrease in HW in the old female GSNOR^-/-^ mice, relative to middle age and significantly relative to their WT counterpart (p=0.02; Figure 3B). The changes in normalized HW correspond to changes in animal size from young to middle-age group of WT and GSNOR^-/-^ mice. Although, the absence of GSNOR appears to reverse this process in the old group of both sexes, the data suggests that in female mice, the absence of GSNOR could affect HW irrespective of animal size.

### GSNOR deletion attenuates age-related cardiac remodeling

To further investigate our observed HW changes, we took a subset of mice and performed conscious serial echocardiography in 3-month intervals starting at 4-months of age. There were striking age-related changes in both male and female WT mice in key echocardiographic parameters (Figure 4): left ventricular relative wall thickness (RWT), calculated based on septal and poster wall diameters, ejection fraction (EF), left ventricular internal diameter in diastole (LVIDd) and left ventricular mass (LV mass). Although the direction of change with age was similar in both sexes in the WT group, there was a sex difference in end point and trajectory, within some parameters. In contrast, age-related changes observed in GSNOR^-/-^ mice were extremely modest and, in some parameters, showed no effect, sex-specific or otherwise.

**Figure 4.**
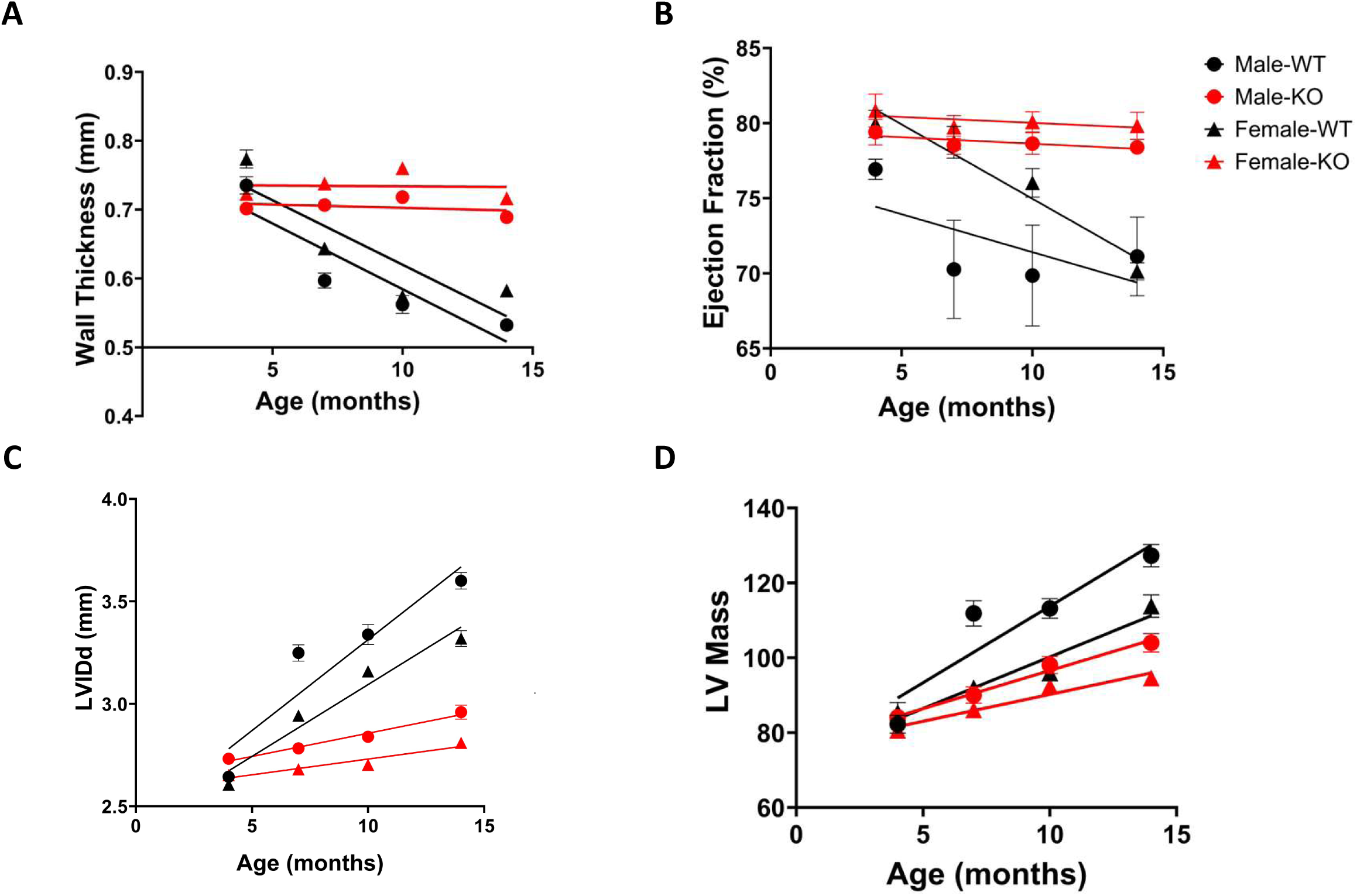
Serial echocardiography measurements of WT and GSNOR-KO mice. (**A**) LV relative wall thickness (**B**) Ejection fraction (**C**) LVIDd and (**D**) LV Mass in male (circle) and female (triangle) WT (black) and GSNOR-KO (red) mice. The effect of age, sex and genotype were determined using a mixed-effects model with Tukey’s multiple comparisons post hoc tests. Simple linear regression was used to determine relationship between each parameter and age. LV- left ventricle; LVIDd- LV internal diameter in diastole; WT-wild-type; KO- S-nitrosoglutathione reductase knockout. Data presents mean±SEM. n=4-10 mice.

RWT was similar at 4 months of age in male and female mice across both genotypes. There was a significant and similar age-related decrease in RWT (p<0.0001; Figure 4A), in both male (r^2^= 0.71) and female (r^2^= 0.67) WT mice. Surprisingly, in GSNOR^-/-^ mice, RWT remained unchanged from 4 to 14 months of age in both male and female mice (Figure 4A). There was no significant deviation from 0 with age in the RWT slopes of both male and female GSNOR^-/-^ mice. The lack of an aging effect on RWT in the GSNOR^-/-^ mice can be explained by the mirroring/opposite pattern observed in both LV posterior wall diameter (Supplementary Figure 1; SF-1) and interventricular septal wall (SF-2) in diastole, relative to WT. Across all four groups, there was a significant interaction between age and genotype in RWT (p<0.001). LVIDd was similar at 4 months of age in male and female mice across both genotypes (Figure 4B). There was a significant increase with age in both male and female WT mice (p<0.0001). Although this pattern of increase was similar across all ages in both males (r^2^= 0.8) and females (r^2^= 0.87), LVIDd was greater in males for all ages post 4 months relative to female mice. In GSNOR^-/-^ mice, LVIDd followed a similar trend in increase from 4 months in both sexes but this significantly attenuated relative to WT (p<0.0001; Figure 4C). There was a strong interaction between age, sex, and genotype (p=0.057).

Furthermore, EF significantly decreased with age in WT female mice (r^2^= 0.63; p<0.001) and trended towards significance in WT males (r^2^=0.04; p=0.18; Figure 4B). While, in GSNOR^-/-^ mice, EF remained unchanged from 4 to 14-months of age (Figure 4B) in both sexes. Across all 4 groups there was a significant interaction between age and genotype (p=0.014). Additionally, LV mass was similar in all groups at 4 months of age (Figure 4D). In WT, as expected, LV mass was greater in males for all ages post 4 months relative to female mice and significantly increased with age in both sexes (p<0.001). In GSNOR^-/-^ mice this age-related increase in LV mass was similar in pattern but stunted in extent. These parameters suggest that with age there is a degree of eccentric hypertrophy occurring in WT hearts, shown by the increasing LVIDd, decreasing RWT and EF. Whilst GSNOR^-/-^ hearts on the other hand may be undergoing concentric remodeling evidenced by no change in RWT, modest increase in LVIDd but maintenance of EF.

### GSNOR deletion alters age-related cardiomyocyte function in a sex-dependent manner

Thus far, our findings indicate that GSNOR modulates age-related cardiac structural and functional remodeling. To gain insight into the role of GSNOR in age-related functional changes on a cellular level, we isolated and paced cardiomyocytes from a subset of WT and GSNOR^-/-^ mice aged 3-4 months and 14-15 months and recorded baseline sarcomere shortening and calcium flux parameters. We chose these two age ranges, young to mid-age because this was the range which showed a significant increase in heart-weight in female GSNOR^-/-^ mice (Figure 3B).

Age had no effect on peak Ca^2+^ transient amplitude in either WT or GSNOR^-/-^ male mice (Figure 5A). The time-to-peak Ca^2+^ was significantly increased in middle-age WT males (p=0.004 SF-4A), but this was not observed in GSNOR^-/-^ males. Additionally, Ca^2+^ relaxation time-to-50% ([Ca^2+^]_BL50_) was significantly increased (p=0.0081; Figure 5B) in male WT mice, supporting age-related diastolic dysfunction previously reported [23]. Interestingly, in male GSNOR^-/-^ mice there was no age-related effect on [Ca^2+^]_BL50_ (Figure 5B). In the female groups, there was a significant increase in peak Ca^2+^ transient amplitude with age in GSNOR^-/-^ females alone (p=0.0114; Figure 5C), while time-to-peak Ca^2+^ was unaffected by age in both WT and GSNOR^-/-^ females (SF-4B). There was a significant increase in [Ca^2+^]_BL50_ with age in both WT (p=0.0075) and GSNOR^-/-^ female mice (p=0.024; Figure 5D). Although there was no effect of age on τ, there was a significant decrease in tau (decay is faster) in young GSNOR^-/-^ females relative to age-matched female WT (p=0.0043; SF-4B). This data indicates that GSNOR modulates age-related functional changes in a sex-specific manner.

**Figure 5.**
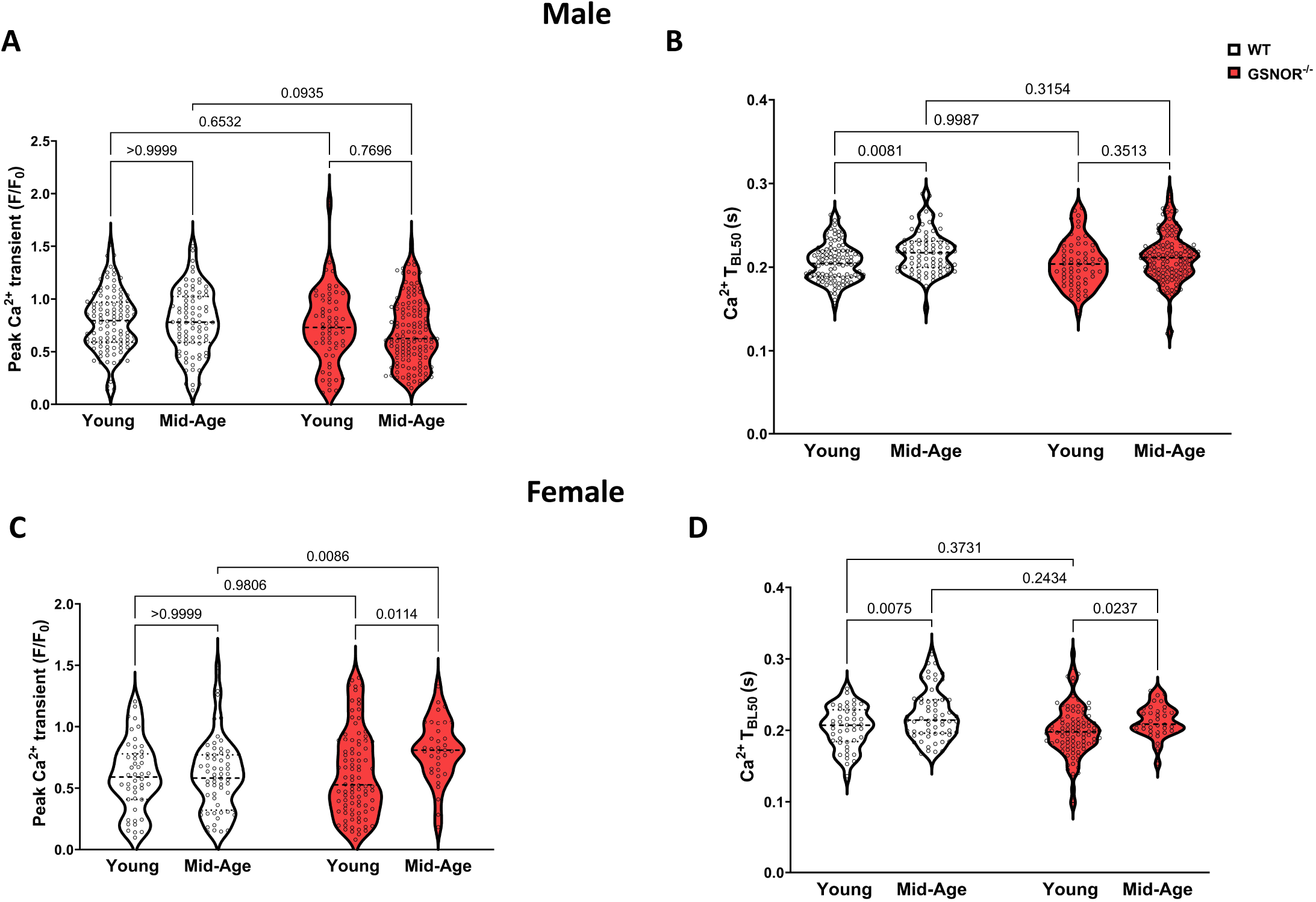
Calcium transients in cardiomyocytes. Quantified recordings of calcium transients of Fura-2AM loaded cardiomyocytes paced at 1Hz isolated from young (3-4 months) and mid-age (13-15 months), WT (grey) and GSNOR-KO (red), male (**A**) and female (**B**) mice. Statistical significance was determined at p<0.05 by two-way ANOVA with Tukey’s post hoc multiple comparisons test. N=36-131 cells; 7-11 mice.

Additionally, we observed a significant decrease in sarcomere shortening with age in male WT mice (p=0.015; Figure 6A). In young GSNOR^-/-^ males sarcomere shortening was significantly decreased, compared to young WT males (p=0.0001). Although there was no effect of age on sarcomere shortening in male GSNOR^-/-^ mice (p=0.834), we did observe a significant increase in time-to-peak shortening with age in male GSNOR^-/-^ mice (p=0.0001; SF-3A) which was not observed in male WT. In female mice, there was no effect of age on sarcomere shortening in the WT group (Figure 6C). Surprisingly, young GSNOR^-/-^ females showed a significant increase in sarcomere shortening relative to young WT females (p=0.0025), but a significant decrease in sarcomere shortening with age (p=0.0019), not observed in WTs. There was a significant increase in time-to-peak shortening with age in female WT mice (p=0.038; SF-3B), not observed in GSNOR^-/-^ mice. Together these data suggest that the absence of GSNOR in males attenuates age-related cardiomyocyte calcium dysfunction, yet in females GSNOR deletion might exacerbate this dysfunction.

**Figure 6.**
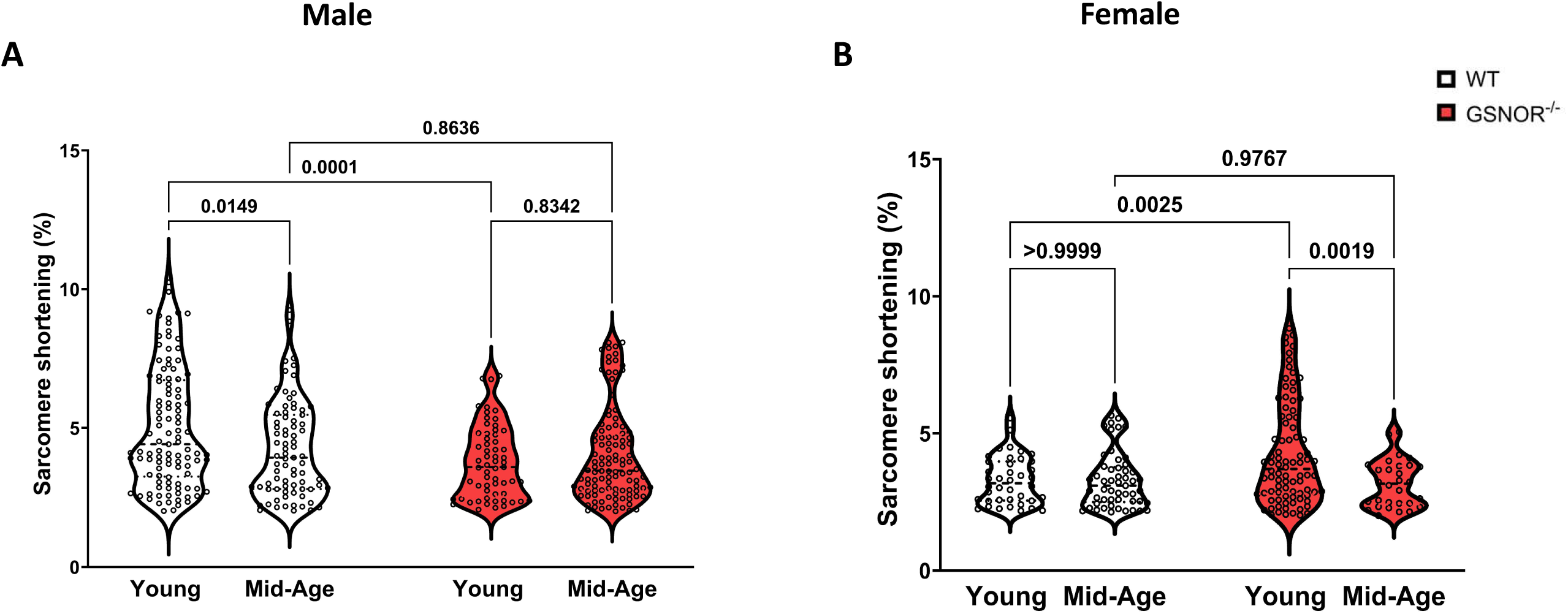
Percentage of sarcomere shortening in cardiomyocytes. Quantified recordings of sarcomere shortening in Fura-2AM loaded cardiomyocytes paced at 1Hz isolated from young (3- 4 months) and mid-age (13-15 months), WT (grey) and GSNOR-KO (red), male (**A**) and female (**B**) mice. Statistical significance was determined at p<0.05 by two-way ANOVA with Tukey’s post hoc multiple comparisons test; n=36-131 cells; 7-11 mice.

### GSNOR deletion alters aging-related cardiac S-nitrosylated protein profile in females

To determine the potential mechanisms through which GSNOR deletion might be exacerbating cardiomyocyte dysfunction in females, we performed SNO-RAC with LC-MS/MS in WT and GSNOR-KO females, to assess potential changes in the SNO proteome. As previously reported a number of the SNO-modified proteins identified were predominantly mitochondrial [19]. We identified 54 common SNO-modified peptides in both young and middle age WT females, whereas, 61 unique peptides were identified in young WT females relative to the 26 unique peptides in the middle-age group (Figure 7A). Of these, two mitochondrial matrix proteins were significantly downregulated in middle-age WT females and therefore SNO-enriched in young WT females, dihydrolipoyl dehydrogenase (DLD; O08749) at Cys80 and Cys85 and the β-subunit of E1 component of pyruvate dehydrogenase (PDHB; Q9D051) at Cys263 relative to middle age counterparts (Figure 7B).

**Figure 7:**
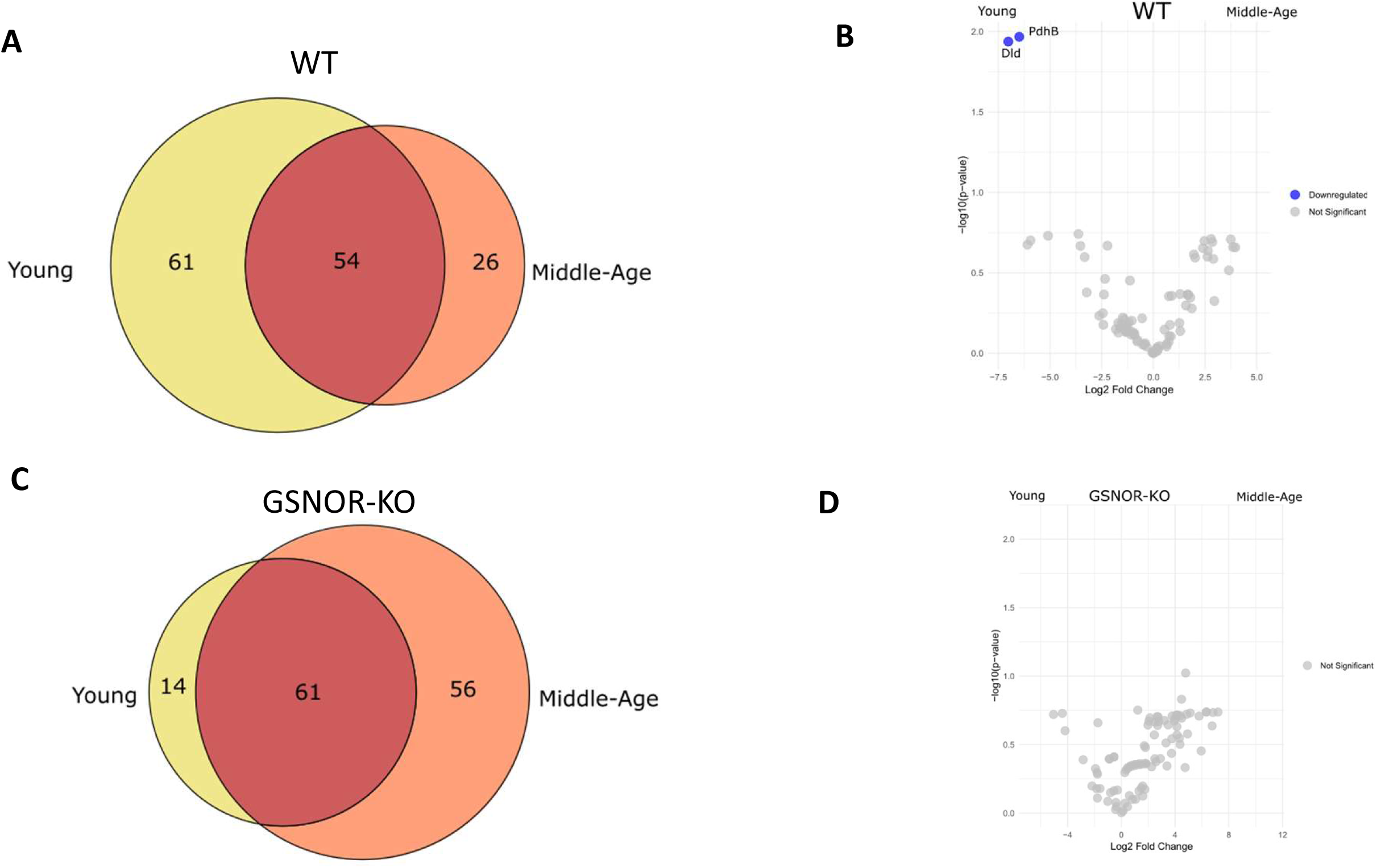
SNO-profile in WT and GSNOR-KO female mice. Venn diagram showing shared and unique SNO-modified peptides in female WT (**A**), young (3-4 months) and middle-age (13-15 months) mice and volcano plot of age-comparison (**B**), upregulated peptides indicated in blue. (**C**) Venn diagram showing shared and unique SNO-modified peptides in GSNOR-KO young and middle-age mice. Comparison between ages shown in volcano plot (**D**). Statistical significance was determined at p<0.05 with Mann-Whitney test for non-parametric data and for parametric data type 2 or 3 Student’s T-test was used, pre-determined by F-test. n=3/group.

In the GSNOR-KO females, 61 common SNO-modified peptides were identified as in both young and middle age mice. Only 14 unique peptides were identified in young GSNOR-KO females, relative to 56 uniquely expressed SNO-modified peptides in middle-age group (Figure 7C). Interestingly, however, no peptide group reached the statistically significant threshold of p<0.05 as being enriched in middle age vs young within female GSNOR-KO mice (Figure 7D). Together these data suggest that in WT females, there is an age-dependent shift in the SNO-status of certain metabolic proteins with age, and the absence of GSNOR removes this imbalance.

## Discussion

In the present study, we investigate the sex dependent effects of GSNOR deficiency in murine cardiac aging, and its impact on maladaptive age-related changes in the heart. To the best of our knowledge this is the only study with data showing that GSNOR deficiency is associated with beneficial maintenance of overall cardiac function with age in both sexes; but key maladaptive molecular and cellular changes, resulting in impaired cardiomyocyte function, occur only in aging females.

Sex and age are both non-modifiable risk factors for cardiovascular disease. Prevalence of adverse cardiac events increases with age, and the rate of this increase is dependent upon sex. In preclinical and human studies female specific cardioprotection has been directly linked to nitric oxide signaling [7]. Protein SNO, regulated by GSNOR activity, has been established by our group as a mechanism by which preconditioning and other cardioprotective interventions confer protection in myocardial I/R injury [4, 6, 24, 25]. Thus, GSNOR inhibition or deletion has been established as advantageous post *ex-vivo* in I/R injury [8, 10], and *in-vivo*, post myocardial infarction (MI) [9, 12]. These observations, however, were made in young male animals only. Moreover, Casin *et al* noted that female-specific cardioprotection in I/R injury was lost with GSNOR inhibition [8]. Female-specific protection from cardiac injury has historically been documented [26, 27] and linked to higher SNO levels and GSNOR activity in the female heart as we have previously reported [8].

Emerging reports suggest that GSNOR also plays a role in aging [11, 28]. We have shown for the first time that GSNOR activity in the heart is inversely proportional to age (Figure 1). Moreover, myocardial GSNOR protein levels in old mice (>18 months) significantly decrease (Figure 2). This age-associated decline in GSNOR activity has previously only been observed in brain homogenates of WT mice [28], but biological sex was not considered. Importantly, we show for the first time that this reduced GSNOR activity with age occurs in females alone, while both GSNOR activity and protein levels are unchanged in males. This supports the notion that cardiac function decline operates by different mechanisms in males v females.

Sex differences in humans and animals are further emphasized with age, in that the increased heart weight is more pronounced in males relative to females (Figure 3),[21, 29], most likely due to the naturally hypertrophic effect of generally higher levels of testosterone in males than females. Remarkably we found that in the absence of GSNOR, this aging effect is attenuated in males, while in females without GSNOR, there appears to be an age-dependent biphasic effect on heart weight. Controversy exists in the literature regarding the role of GSNOR in the acceleration or deceleration of the aging process. Salerno *et al* [30] showed that inhibition of GSNOR accelerated differentiation and maturation of inducible pluripotent stem cells, suggesting that GSNOR slows down maturation and thus aging. In the brain and liver, Rizza *et al* [28] showed that GSNOR deficiency promoted mitophagy, a feature of cardiac aging, thus supporting the idea of GSNOR as a promoter of longevity. Rizza *et al* [28] further reinforced this notion by showing in their report that peripheral blood mononuclear cells from healthy long-lived individuals (greater than 95 years of age) maintained high GSNOR mRNA levels. These findings, however, are in contrast with a report by Zhang *et al* [11] wherein GSNOR overexpression in transgenic (TG) mice resulted in cognitive decline. The report concludes by presenting GSNOR as a potential target for the treatment of age-related cognitive impairment. We, however, suggest that there is a degree of sex-specificity that was unaccounted for in the Zhang *et al* study. Zhang *et al* [11] report that male and female mice were included in their TG experiments. Although the overall finding of cognitive decline was observed in both sexes, the pattern differed between the sexes. Additionally, the rescue experiments in that study were done on only male GSNOR-KO mice [11].Our findings, taken together with the above literature suggests that the balance of GSNOR activity might be the key to maintaining health in either sex.

Our observations suggest that GSNOR deletion decreases the rate of cardiac aging by maintaining EF and ventricular wall thickness in both sexes (Figure 4). However, when taken together with cardiomyocyte functional data, this decrease in cardiac aging appears to be male-specific. In males, sarcomere shortening, and calcium transients are both maintained (Figures 5- 6). While in females, the opposite is observed; particularly from young to middle-age (Figures 5- 6), GSNOR deletion appears to accelerate cardiac aging, or at least exacerbate it. This finding is supported by a previous report in which impaired cardiomyocyte β-adrenergic responsiveness, a characteristic of aging, was documented in female GSNOR^-/-^ mice alone [31]. This observation of GSNOR as a potential genetic determinant of outcome in females was first observed in LPS-challenged mice [32], where LPS-challenged female GSNOR^-/-^ mice experienced a 10-fold increase in mortality vs. WT, compared with the 2-fold observed in males [32].

In our SNO-RAC experiments, we found that in the female WT group, key mitochondrial proteins DLD and PDHB were identified as SNO-modified in young females, but not in middle-age (Figure 7), which is interesting given that we find GSNOR activity to be decreased in this timeframe. Nonetheless, both proteins are components of the pyruvate dehydrogenase complex (PDC) [33] specifically converting pyruvate to acetyl-CoA and are involved in glucose metabolism. Interestingly PDHB has previously been reported to be involved in myogenic differentiation and expression levels have been shown to decrease in skeletal muscle with aging [34]. As such, a crucial follow up study investigating how denitrosylation of PDHB modulates pyruvate dehydrogenase complex activity and its direct impact on cardiac function is encouraged. Equally important would be to specifically target and follow the SNO-state of certain crucial protein markers previously implicated in cardiac aging and senescence. Data from these experiments could aid in providing a direct cause and effect relationship between changes in the SNO state of key proteins and cardiac phenotypic changes. Additionally, in female GSNOR^-/-^ mice, we did not find any SNO proteins that were significantly changed with age between experimental groups, which is noteworthy considering that GSNOR acts as a denitrosylase. In any case, the overall SNO proteome did appear to shift differently in WT vs. GSNOR^-/-^ female hearts with aging.

A number of limitations of this study require acknowledgement. Firstly, our study examined male and female WT and GSNOR^-/-^ mice at young and middle-age, but our old-age timepoints are limited. Thus, future studies will examine extended timepoints in aging. Secondly, we measured cardiomyocyte function at baseline in cardiomyocytes, albeit paced at low frequency (1Hz). Future studies will determine the response to various substrates mimicking endogenous activation such as isoproterenol and fatty acids at the same aging timepoints, and additionally, determine whether at an older age (>20months) the response of cardiomyocytes from both male and female GSNOR-KO mice eventually converge. We are also limited in drawing conclusions on sex-specific differences in SNO-modification due to our focus on female mice in the SNO-RAC experiments. As such, we encourage future work to investigate the myocardial proteome in male mice to explore the changes within their SNO profile with age.

In summary, our data highlights the significance of GSNOR in the sex disparity of cardiac aging. On the surface, GSNOR deficiency appears beneficial to cardiac aging in both sexes, yet at the cellular level we have revealed a sex-disparity and the potential for underlying cellular dysfunction, particularly in female hearts. Taken together, GSNOR deficiency may present a mechanism through which the female heart specifically is at a higher risk of age-related CVD and may represent a potential clinical target.

---

**SF1:**
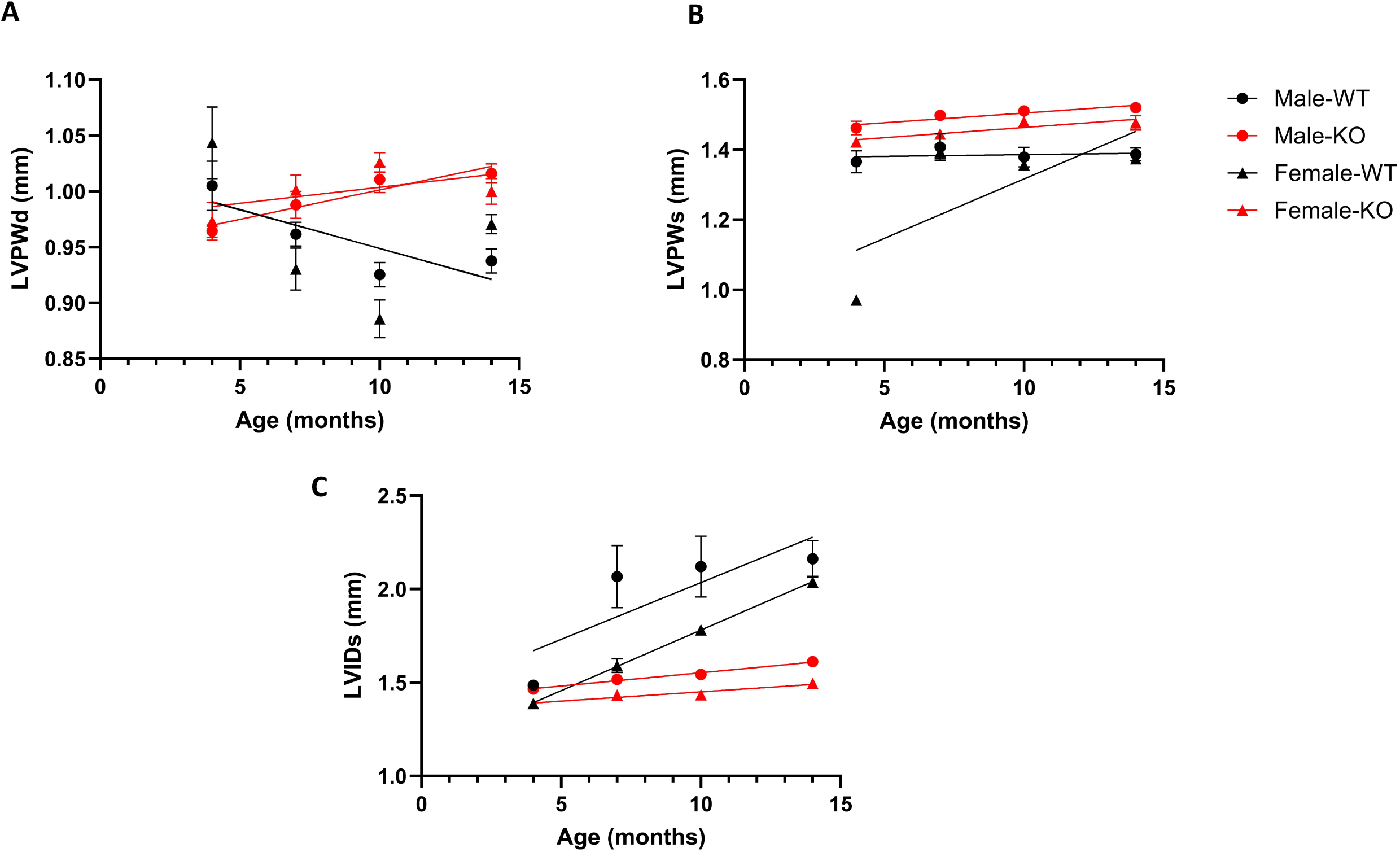
Echocardiography parameters in Male and Female WT and GSNOR-KO mice. A.) Left ventricular posterior wall in diastole (LVPWd). B) Left ventricular posterior wall in systole (LVPWs) C.) Left ventricular interior diameter systole (LVIDs) in male (circle) and female (triangle) WT (black) and GSNOR-KO (red) mice. Simple linear regression was used to determine relationship between each parameter and age. LV- left ventricle; LVIDd- LV internal diameter in diastole; WT-wild-type; KO- S-nitrosoglutathione reductase knockout. Data presents mean±SEM. n=4-10 mice.

**SF2:**
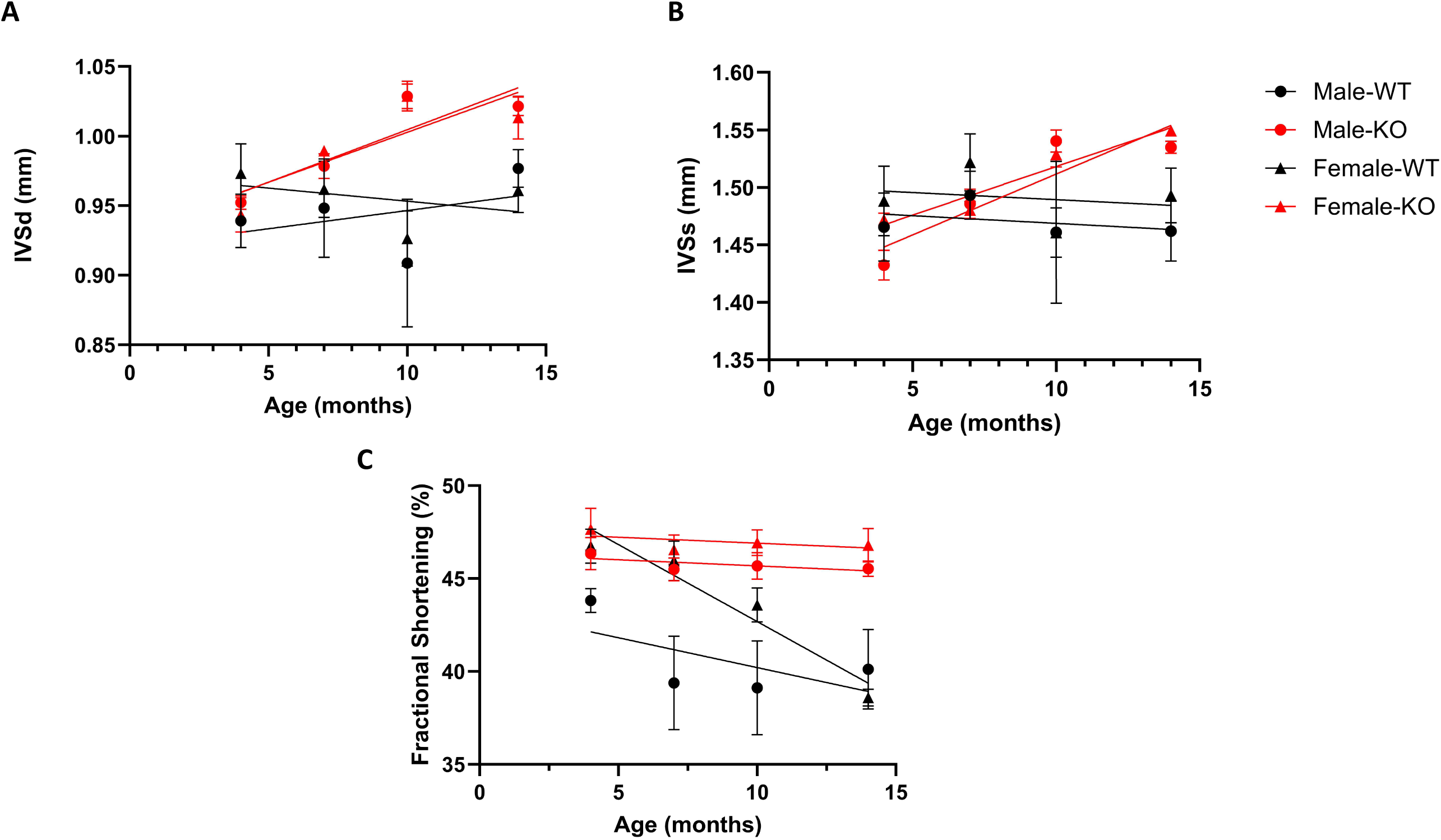
Echocardiography parameters in Male and Female WT and GSNOR-KO mice. A.) Internal ventricular septum in diastole (IVSd). B) Internal ventricular septum in systole (IVSs) C.) Fractional Shortening in male (circle) and female (triangle) WT (black) and GSNOR-KO (red) mice. Simple linear regression was used to determine relationship between each parameter and age. LV- left ventricle; LVIDd- LV internal diameter in diastole; WT-wild-type; KO- S-nitrosoglutathione reductase knockout. Data presents mean±SEM. n=4-10 mice.

**SF3:**
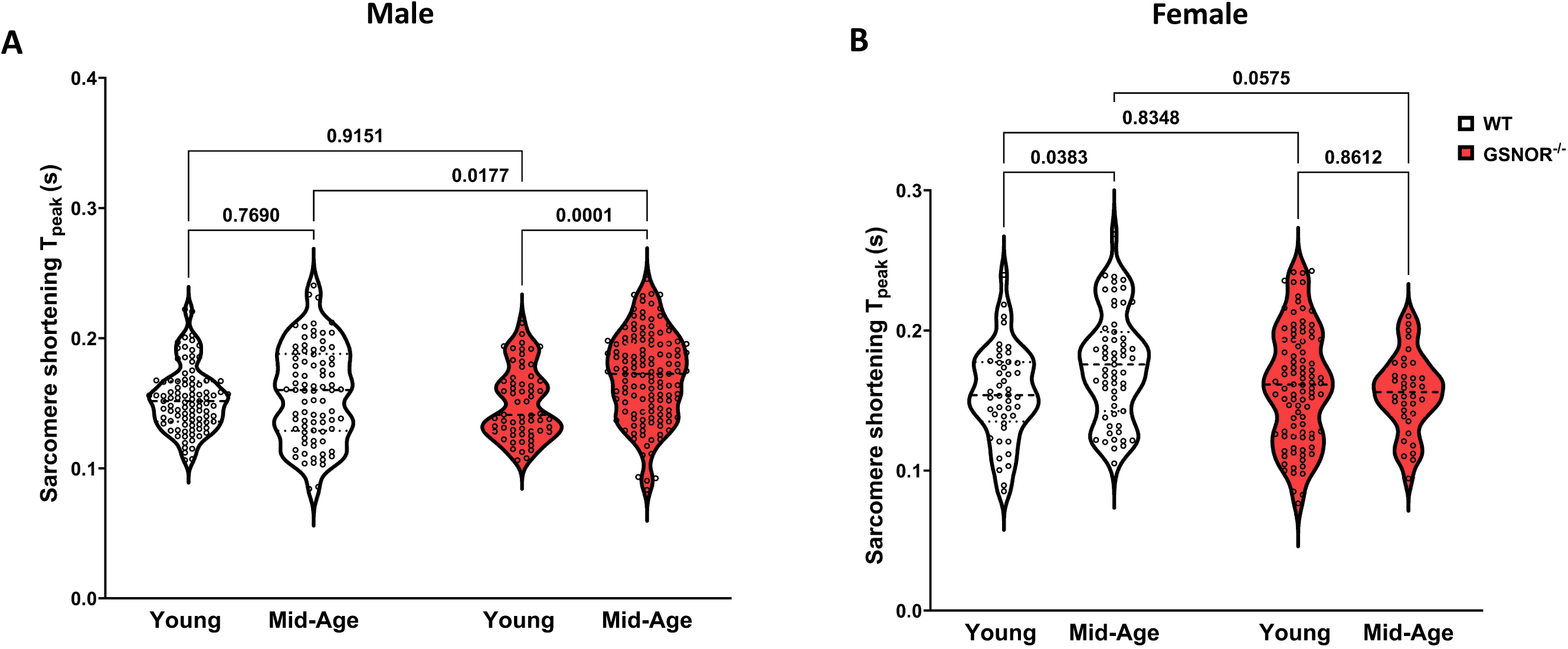
Sarcomere shortening kinetics in cardiomyocytes. Quantified recordings of the time to peak sarcomere shortening of Fura-2AM loaded cardiomyocytes paced at 1Hz isolated from young (3-4 months) and mid-age (13-15 months), WT (grey) and GSNOR-KO (red), male (**A**) and female (**B**) mice. Statistical significance was determined at p<0.05 by two-way ANOVA with Tukey’s post hoc multiple comparisons test and n=36-131 cells; 7-11 mice.

**SF4:**
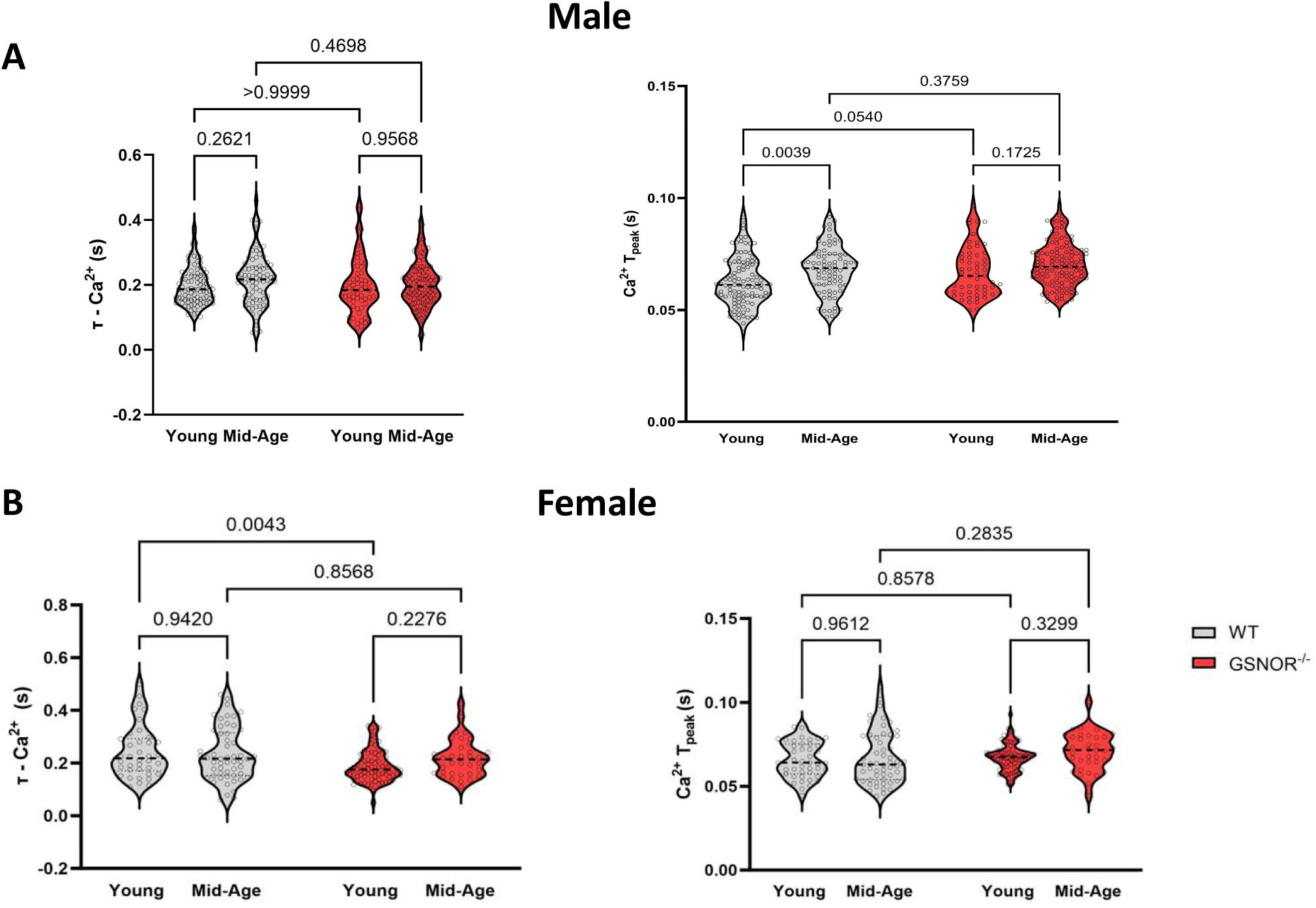
Calcium kinetics parameters in cardiomyocytes. Quantified recordings of calcium relaxation (tau) and time to peak of Fura-2AM loaded cardiomyocytes paced at 1Hz isolated from young (3-4 months) and mid-age (13-15 months), WT (grey) and GSNOR-KO (red), male (**A**) and female (**B**) mice. Statistical significance was determined at p<0.05 by two-way ANOVA with Tukey’s post hoc multiple comparisons test. N=36-131 cells; 7-11 mice.

## Notes

### Competing Interest Statement

The authors have declared no competing interest.

### Funding

- National Institutes of Health: T32HL007227, T32ES007141, F31HL165820, R21HL157800, R01HL136496

